# β-Hydroxybutyrate elicits divergent metabolic responses between MCF-7 and T47D ER+ breast cancer cells under glucose restriction

**DOI:** 10.64898/2026.05.14.725288

**Authors:** Cynthia Cheung, Natalija Glibetic, Rylee Maldonado, Scott Bowman, Tia Skaggs, Louise Torres, Katelynn A. Perrault Uptmor, Michael Weichhaus

## Abstract

**Background:** The ketogenic diet is being explored as an adjuvant intervention in breast cancer because it lowers circulating glucose and elevates ketone bodies such as β-hydroxybutyrate (BHB), but how individual ER+ breast cancer subtypes adapt to these conditions remains poorly characterized. We examined metabolic responses to BHB supplementation under glucose restriction in two ER+ breast cancer cell lines, asking whether metabolic adaptation patterns differ between models.

**Methods:** MCF-7 and T47D cells were cultured under high glucose, glucose-restricted (5% of standard), or glucose-restricted with 10 mM BHB conditions and profiled by comprehensive two-dimensional gas chromatography–mass spectrometry (GC×GC-MS). Pairwise Welch’s t-tests with Benjamini–Hochberg false discovery rate (FDR) correction were applied to identify treatment-responsive metabolites. Targeted assays quantified intracellular glycine, SHMT1 protein, and total branched-chain amino acid (BCAA) concentrations across a BHB dose range (2.5-15 mM). Patient tumor transcriptomic data from TCGA (n=1,084) and paired tumor-normal samples from GSE58135 (n=20) were analyzed for genes involved in one-carbon, ketone body, and BCAA metabolism.

**Results:** MCF-7 and T47D cells exhibited markedly divergent metabolic responses to BHB. In MCF-7 cells, BHB supplementation produced a broad pattern-level metabolic shift: 75% of detected metabolites trended upward when BHB was added to glucose-restricted cultures (C vs. B comparison), with 1,4-butanediol reaching nominal significance (FC=2.35, p=0.016) and a 4.1-fold trend increase in lactic acid (p=0.11), although no individual metabolite survived FDR correction. T47D cells showed essentially no metabolic response to BHB at the global level. Targeted assays detected an elevation in glycine at 5 mM BHB in both cell lines that did not follow a monotonic dose response and was not accompanied by changes in SHMT1 protein expression. Total BCAA levels were elevated by BHB in T47D cells but remained unchanged in MCF-7 cells. In paired patient samples, OXCT1 (log2FC = −1.41), SHMT1 (log2FC = −1.31), and ACAT1 (log2FC = −1.07) were significantly downregulated in ER+ tumors relative to matched normal tissue (adjusted p < 0.001 for all three).

**Conclusions:** ER+ breast cancer cell lines show heterogeneous metabolic responses to BHB supplementation under glucose restriction. The broad pattern of metabolite elevation in MCF-7 but not T47D cells suggests that capacity to utilize ketone bodies as metabolic substrate varies between ER+ models. The downregulation of OXCT1, ACAT1, and SHMT1 in ER+ tumors compared to normal tissue identifies these enzymes as candidate biomarkers that may help stratify which patients are likely to benefit from ketogenic interventions. Findings related to individual metabolites should be regarded as exploratory and require validation in larger, adequately powered cohorts.

## 1. Background

Breast cancer remains one of the most prevalent malignancies affecting women worldwide, accounting for approximately 2.3 million new cases and over 685,000 deaths annually [1]. Estrogen receptor-positive (ER+) subtypes comprise approximately 70–80% of cases, making them the most prevalent form. While endocrine therapies targeting ER signaling have significantly improved outcomes, resistance develops in up to 50% of advanced cases, often driven by metabolic adaptations that sustain tumor growth despite hormone deprivation [2,3]. Research suggests ER+ breast cancer cells exhibit heightened glucose dependency via the Warburg effect, where aerobic glycolysis supports proliferation and survival, potentially rendering them more susceptible to metabolic interventions [4,5].

Studies indicate that ER signaling directly reprograms metabolism in ER+ cells, upregulating glycolytic enzymes and altering pathways such as choline and amino acid metabolism, which contribute to endocrine resistance [6]. Glucose metabolic reprogramming has been linked to acquired resistance to CDK4/6 inhibitors in ER+ models, with sensitive cells showing enhanced aerobic glycolysis that shifts upon resistance [7]. Metabolic stresses such as serine or glucose deprivation can silence ERα expression through epigenetic mechanisms, highlighting the interplay between nutrient availability and hormone signaling in ER+ tumors [8].

The ketogenic diet (KD), defined by its high-fat (typically 70–80%), moderate-protein, and very low-carbohydrate composition, induces a state of nutritional ketosis where the liver produces ketone bodies such as β-hydroxybutyrate (BHB) as alternative energy substrates, effectively mimicking fasting and reducing the serum glucose and insulin levels that cancer cells exploit for growth [9]. Preclinical and clinical studies suggest that KD may inhibit tumor proliferation, enhance chemotherapy sensitivity, and improve patient outcomes in breast cancer by targeting glycolytic pathways, though evidence remains mixed and is often limited to animal models or small trials [10,11,12]. The biological effects of BHB itself are similarly heterogeneous: in some contexts BHB reduces cancer cell viability and induces apoptosis [13,14], whereas in others it promotes chemoresistance and proliferation in breast cancer cells [15].

Amino acid metabolic reprogramming plays crucial roles in cancer cell survival and proliferation. Branched-chain amino acids (BCAAs), valine, leucine, and isoleucine, contribute significantly to cellular energy production in breast cancer, with metabolic flux analysis revealing that 67% of carbon in mevalonate synthesis derives from leucine degradation in MCF-7 cells [16,17]. One-carbon metabolism pathways, particularly involving serine hydroxymethyltransferase (SHMT) enzymes, are frequently dysregulated in breast cancer [18]. SHMT2 expression correlates with tumor grade and poor prognosis in breast cancer patients [19], whereas its cytosolic counterpart SHMT1 catalyzes the reversible conversion of serine to glycine and contributes to nucleotide biosynthesis and redox homeostasis [20,21]. Importantly, serine rather than glycine serves as the primary one-carbon donor for cancer cell proliferation, as exogenous glycine cannot replace serine to support nucleotide synthesis [22].

Despite these findings, the metabolic adaptations of breast cancer cells to ketone body supplementation under glucose-scarce conditions remain incompletely characterized, and the extent to which different ER+ cell line models share or diverge in their responses is unclear. Cell line heterogeneity within ER+ breast cancer is well documented at the proteomic and bioenergetic level, with MCF-7 cells relying more heavily on oxidative phosphorylation than T47D cells, which display higher glycolytic flux and greater proton leak [23,24]. Whether such differences extend to ketone body utilization has not been systematically examined.

In the present study, we addressed these gaps using paired MCF-7 and T47D models exposed to glucose deprivation with BHB supplementation. We performed comprehensive GC×GC-MS metabolomics with FDR-controlled statistical analysis, complemented by targeted quantification of glycine, SHMT1, and total BCAAs across a BHB concentration range. To place our in vitro observations in a clinical context, we examined co-expression patterns between one-carbon, BCAA, and ketone body metabolism genes in the TCGA breast cancer cohort and performed paired differential expression analysis in ER+ tumor and matched normal tissue from the GSE58135 dataset. The integrated analysis emphasizes cell line heterogeneity as a primary feature of the metabolic response to BHB and identifies a coordinated set of ketone body catabolic enzymes that are downregulated in ER+ tumors.

## 2. Methods

### 2.1 Cell culture

Human breast cancer cell lines MCF-7 (Cat No.: HTB-22) and T47D (Cat No.: HTB-133) were purchased from the American Tissue Culture Collection (ATCC, Manassas, VA, USA) and routinely maintained in DMEM (Gibco, ThermoFisher, Waltham, MA, USA) with 10% FBS (R&D Systems, Minneapolis, MN, USA), 2 mM glutamine (Gibco) and 50 ng/mL gentamicin (Lonza Biologicals, Portsmouth, NH, USA). Cell lines were incubated at 37 °C in humidified 5% CO₂-supplemented air and sub-cultured at approximately 80% confluence. Cell cultures were inspected daily for visible contamination and consistent growth rates. The cell culture facility is accessible only to qualified researchers and does not maintain HeLa cells. Experiments were performed within 25 passages of receipt from ATCC.

### 2.2 Sample preparation for metabolomic analysis

In 10 cm dishes, 1×10⁶ MCF-7 or T47D breast cancer cells were incubated in regular DMEM medium or treated with 225 mg/L (5% of glucose in regular DMEM medium) alone or in combination with 10 mM β-hydroxybutyrate (BHB) (MilliporeSigma, St. Louis, MO, USA) for four days, across multiple independent experimental sets. Cells were harvested using 0.25% trypsin (Gibco), pelleted by centrifugation, and stored at −80 °C.

### 2.3 Sample derivatization

Frozen cell pellets were thawed on ice and resuspended in 1 mL of ice-cold extraction buffer consisting of acetonitrile:isopropanol:water (3:3:2, v/v/v). Samples were vortexed five times for 15 seconds each, then frozen in crushed dry ice for 20 minutes. After thawing on ice for 10 minutes, the vortex-freeze-thaw cycle was repeated two additional times. Following extraction, samples were dried by centrifugal evaporation. For derivatization, dried samples were reconstituted with 10 µL of methoxyamine hydrochloride (MOX) reagent (Thermo Fisher Scientific, TS-45950) and incubated at 30 °C for 90 minutes with shaking at 1100 rpm. Subsequently, 90 µL of N-methyl-N-(trimethylsilyl)trifluoroacetamide (MSTFA; Sigma-Aldrich, 394866) was added, and samples were incubated at 37 °C for 30 minutes with shaking at 1000 rpm. Samples were centrifuged at 14,000 rpm at 4 °C, and the supernatant was transferred to GC autosampler vials.

### 2.4 GC×GC-qMS/FID analysis

Samples were analyzed using comprehensive two-dimensional gas chromatography coupled with dual-channel detection by flame ionization detection (FID) and quadrupole mass spectrometry (qMS) on a Trace 1300 GC/FID and ISQ 7000 Single Quadrupole Mass Spectrometer (Thermo Scientific). The first dimension column was an Rxi-624Sil MS column (30 m × 0.25 mm ID × 1.4 µm film, Restek Corporation, Bellefonte, PA, USA); the column junction was equipped with a reverse fill/flush (RFF) INSIGHT flow modulator (SepSolve Analytical Ltd., Peterborough, UK); the second dimension column was a Stabilwax column (5 m × 0.25 mm × 0.25 µm film, Restek). Ultra high purity helium was the carrier gas. The GC oven started at 60 °C, was held for three minutes, ramped at 5 °C/min to 250 °C, and held for five minutes (total run time 46 min). The qMS was operated in electron ionization mode with a scan range of 40–300 amu, scan time of 0.02 s, and overall scan rate of approximately 41.5 scans/s. FID data collection was at 120 Hz. The modulation period was 2.5 s with a 100 ms flush time.

### 2.5 Data processing

Data were acquired using Chromeleon V.7.2.9 (Thermo Scientific) and exported as *.cdf files for processing in ChromSpace V.1.4 (SepSolve Analytical Ltd). Dynamic baseline correction was applied to GC×GC-qMS data (peak width 0.4 s), with peaks integrated using 3-point Gaussian smoothing and minimum height of 300,000. For FID data, Top Hat baseline correction was applied with peak detection minimum height of 1 to accommodate FID sensitivity. Solvent peaks (toluene, hexane), derivatization byproducts (MSTFA-related compounds), and contaminants (butylated hydroxytoluene, plasticizers) were excluded. Identifications were level 2 putative identifications according to the Metabolomics Standards Initiative [25] based on mass spectral library matching and dual retention time data. When multiple peaks corresponded to the same metabolite (e.g., different TMS derivatives), peak areas were averaged.

### 2.6 Metabolomics statistical analysis

Following quality filtering and exclusion of solvent peaks, derivatization byproducts, and contaminants, named metabolites were retained for pairwise statistical comparison; the number of metabolites available varied by comparison according to detection across all samples in both groups being compared. For MCF-7, 46 metabolites were tested in the low glucose versus high glucose comparison (B vs. A), 23 in the BHB-supplemented versus high glucose comparison (C vs. A), and 24 in the primary BHB versus glucose-deprived comparison (C vs. B). For T47D, 24 metabolites were tested in all three pairwise comparisons. Full metabolite-level results, fold changes, raw and adjusted p-values, and coefficients of variation are reported in Supplementary Table S1. Because preliminary inspection revealed substantial inter-day batch effects in the MCF-7 dataset, analyses for MCF-7 were performed on samples from sets 3, 4, and 5, which were all run within a single GC×GC-MS session (n=8 per group) to eliminate inter-day confounding. T47D samples spanned multiple run days (n=14–15 per group) and were analyzed across all available sets. Pairwise comparisons between treatment groups were performed using Welch’s t-tests (unequal variance), and p-values were adjusted for multiple comparisons using the Benjamini–Hochberg false discovery rate (FDR) procedure. Significance thresholds were FDR-adjusted p < 0.05 with |fold change| > 1.5 for strict significance and nominal p < 0.05 for exploratory significance. Principal component analysis (PCA) was conducted on log₂-transformed, standardized data (mean-centered and scaled to unit variance) to visualize treatment group separation. Heatmaps used row-wise z-score normalization with hierarchical clustering. All metabolomics statistical analyses were performed in Python 3.12 with SciPy 1.11 and NumPy.

### 2.7 Glycine assay

In 10 cm dishes, 1×10⁶ MCF-7 or T47D cells were incubated in regular DMEM medium or treated with 225 mg/L glucose (5% of regular DMEM) alone or in combination with 2.5, 5, 10, or 15 mM BHB for four days in three independent experiments. Cells were harvested by 0.25% trypsin, washed three times in ice-cold PBS, resuspended in PBS, sonicated for 15 seconds on ice, and supernatants stored at −80 °C. The Glycine Assay (Cell Biolabs, Cat No. MET-5070, San Diego, USA) was performed in 96-well black microtiter plates per manufacturer’s protocol. Standards were prepared by serial dilution of 10 mM glycine to generate a 0–100 µM standard curve. Fluorescence was measured using a microplate reader (BioRad Mark, Hercules, USA) with excitation at 530–570 nm and emission at 590–600 nm. Samples from three experiments and standards were run in duplicate.

### 2.8 SHMT1 ELISA

Cells were cultured as described in section 2.7. Cell layers were washed in PBS and collected in 1× Cell Lysis Buffer (Cat No. 9803S, Cell Signaling Technologies) with 1× protease inhibitor cocktail (Cat No. P8340, Sigma-Aldrich). SHMT1 levels were quantified using a sandwich ELISA kit (Biomatik, EKU11756). Standards were reconstituted to 5,000 pg/mL and serially diluted to generate a 78–5,000 pg/mL standard curve. Plates pre-coated with anti-SHMT1 antibody were incubated with samples or standards for 1 hour at 37 °C, followed by biotin-conjugated detection antibody (1 hour, 37 °C), avidin-HRP conjugate (30 minutes, 37 °C), and TMB substrate (10–20 minutes, 37 °C protected from light). The reaction was terminated with stop solution, and absorbance was measured at 450 nm. Samples and standards were run in duplicate across three independent experiments.

### 2.9 BCAA assay

Total branched chain amino acid (BCAA) levels were measured using a colorimetric assay kit (Cell Biolabs, Inc., MET-5056). Cells were cultured as described in section 2.7. The reaction mix was prepared by diluting WST-1 reagent 1:10 and NAD⁺ 1:100 in PBS. Standards were prepared by serial dilution of 100 mM L-leucine to generate a 15.6–1000 µM standard curve. Each sample was assayed in paired wells: one positive well containing sample, reaction mix, and leucine dehydrogenase solution, and one endogenous control well lacking the enzyme. After 5–30 minutes of incubation at room temperature on an orbital shaker, absorbance was measured at 450 nm. BCAA concentrations were calculated by subtracting endogenous control values from positive well values and comparing to the leucine standard curve. Three independent experiments were performed with triplicate technical replicates.

### 2.10 Co-expression analysis in TCGA

Gene expression data were obtained from cBioPortal (https://www.cbioportal.org). The Breast Invasive Carcinoma (TCGA, PanCancer Atlas) dataset (1,084 samples) was queried for three primary one-carbon and BCAA metabolism genes (SHMT1, SHMT2, BCAT1) and three ketone body metabolism genes (OXCT1, ACAT1, BDH1). mRNA expression data were analyzed using z-scores relative to normal samples (log RNA Seq V2 RSEM). Both Pearson and Spearman correlation coefficients with corresponding p-values were calculated for each gene pair. Only correlations reaching statistical significance after Bonferroni correction across the nine tested pairs (p < 0.0056) and with |r| > 0.10 are reported in Figure 4; the full correlation matrix is presented in Supplementary Table S2.

### 2.11 RNA-seq differential expression analysis

Raw RNA-seq count data for GSE58135 [26] were obtained from the Gene Expression Omnibus using the NCBI-generated raw counts matrix (GRCh38.p13 annotation). ER+ breast cancer primary tumor samples and their matched adjacent uninvolved breast tissue samples were selected, and strict 1:1 matching was applied to retain patients with exactly one tumor and one matched normal sample, yielding 20 paired samples. Genes with fewer than 10 counts in at least 2 samples were excluded. Differential expression analysis was performed using DESeq2 (v1.48.1) in R (v4.5.1) with a paired study design (∼patient + condition). Log₂ fold changes and Benjamini–Hochberg-adjusted p-values were calculated. Analysis focused on nine genes involved in ketone body, one-carbon, and BCAA metabolism: BDH1, BDH2, OXCT1, OXCT2, ACAT1, ACAT2, SHMT1, SHMT2, and BCAT1. Expression values were variance-stabilizing transformed (VST) for visualization.

### 2.12 Statistical analysis of targeted assays

Glycine, SHMT1, and BCAA assay data were analyzed using GraphPad Prism (version 10.5, GraphPad Software, San Diego, CA). Experiments were performed in biological triplicate. Six experimental groups were compared: control (untreated cells in standard glucose medium), glucose restriction (5% of standard glucose), and four BHB treatment groups (2.5, 5, 10, and 15 mM BHB) in glucose-restricted medium. Data are presented as mean ± standard deviation (SD) or standard error of the mean (SEM) as indicated. Normal distribution was assessed using the Shapiro-Wilk test. For multiple group comparisons, one-way analysis of variance (ANOVA) was performed, followed by Tukey’s multiple comparisons post-hoc test. P < 0.05 was considered statistically significant: *P < 0.05, **P < 0.01, ***P < 0.001, ****P < 0.0001.

## 3. Results

### 3.1 GC×GC-MS metabolomics reveals cell line-specific responses to BHB under glucose restriction

To characterize the metabolic adaptations of ER+ breast cancer cells to glucose deprivation and ketone body supplementation, we performed GC×GC-MS metabolomics profiling of MCF-7 and T47D cells cultured under high glucose, low glucose (5% of standard), or low glucose supplemented with 10 mM BHB. After quality filtering and exclusion of solvent peaks, derivatization byproducts, and contaminants, named metabolites were retained for pairwise comparison. Because metabolite detection varies across treatment conditions, the number of metabolites available for analysis differed by comparison: in MCF-7, 46 metabolites were tested for low glucose vs. high glucose, 23 for BHB-supplemented vs. high glucose, and 24 for the primary BHB vs. glucose-deprived comparison; in T47D, 24 metabolites were tested in all three comparisons (Supplementary Table S1). We applied pairwise Welch’s t-tests with Benjamini–Hochberg FDR correction to identify treatment-responsive metabolites (Fig. 1; Supplementary Table S1).

**Figure 1.**
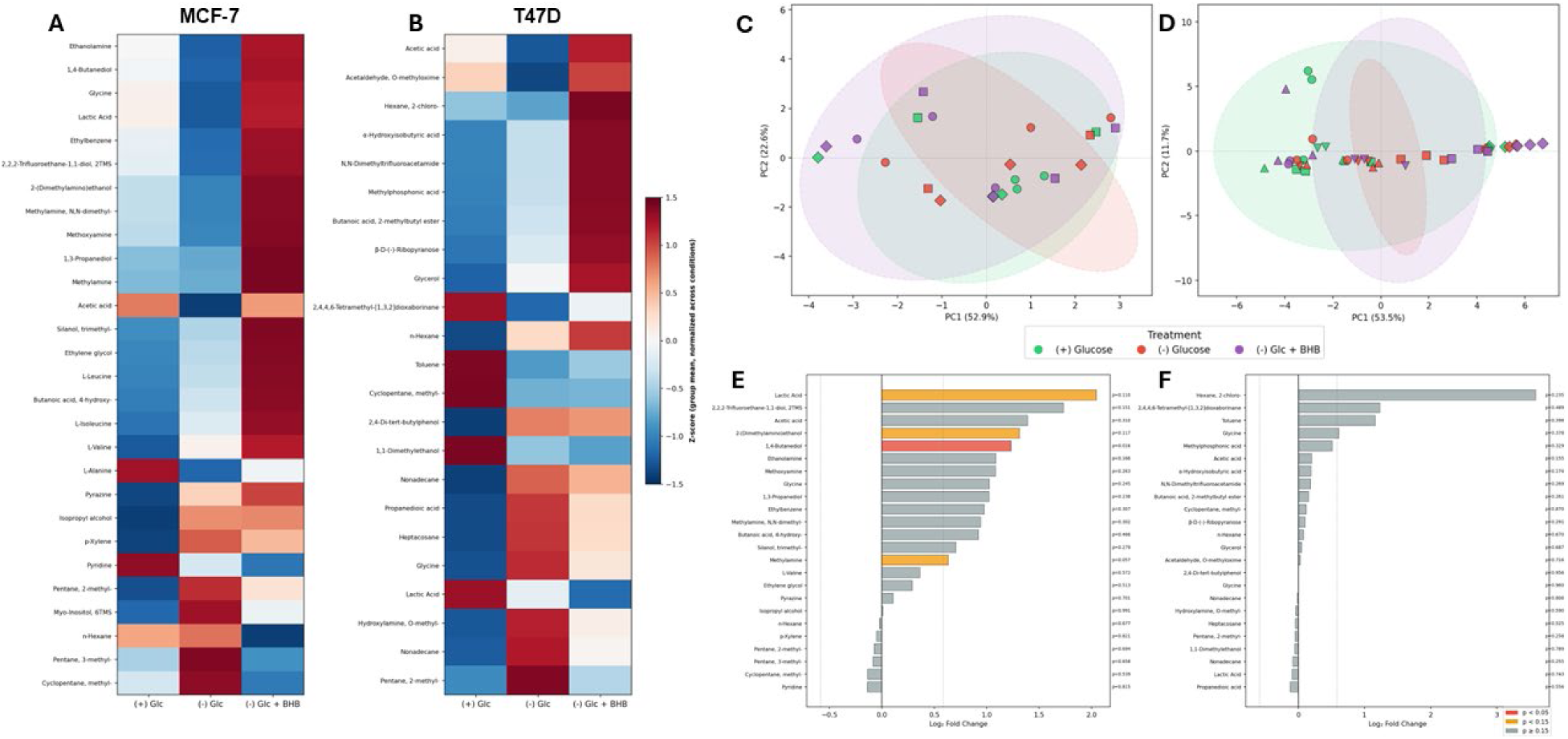
GC×GC-MS metabolomics of glucose-restricted breast cancer cells with β-hydroxybutyrate (BHB) supplementation. MCF-7 and T47D cells were cultured under high glucose (standard DMEM), low glucose (5% of standard), or low glucose supplemented with 10 mM BHB for four days. (A, B) Heatmaps displaying z-score normalized group-mean metabolite abundance for MCF-7 (A) and T47D (B) cells; rows are individual metabolites ordered by clustering, columns are treatment groups. Color scale indicates relative abundance (red: elevated; blue: reduced). (C, D) PCA scores plots for MCF-7 (C) and T47D (D) showing sample distribution by treatment: high glucose (green), low glucose (red), and low glucose + BHB (purple). Variance explained by PC1 and PC2 is indicated on each axis. (E, F) Log₂ fold-change bar charts for the BHB vs. low-glucose comparison in MCF-7 (E) and T47D (F), with p-values from Welch’s t-tests annotated; bars colored by significance level (red: p < 0.05; orange: p < 0.10; yellow: p < 0.15). Analysis applied Benjamini–Hochberg FDR correction; no metabolites reached FDR-adjusted significance, and all metabolite-specific findings are reported as exploratory.

Heatmap visualization of z-score normalized metabolite abundance revealed visually striking but cell line-specific responses to BHB (Fig. 1A, B). In MCF-7 cells (Fig. 1A), the addition of BHB to glucose-restricted cultures produced a coordinated upward shift across the majority of detected metabolites, with ethanolamine, 1,4-butanediol, glycine, lactic acid, ethylbenzene, and 2-(dimethylamino)ethanol among the metabolites showing the most pronounced increases relative to glucose deprivation alone. In contrast, T47D cells (Fig. 1B) displayed a more variable response with no clear coordinated direction across the metabolite panel, although a subset of metabolites including acetaldehyde, hexane derivatives, and α-hydroxyisobutyric acid showed treatment-associated variation.

Principal component analysis (PCA) reflected this difference (Fig. 1C, D). In MCF-7 cells, PC1 and PC2 captured 52.9% and 22.6% of the total variance respectively; the three treatment groups overlapped substantially in score space, consistent with high within-group biological variability, but BHB-treated samples (purple) tended toward more negative PC2 values, indicating a directional treatment effect within a noisy baseline. In T47D cells, PC1 explained 53.5% and PC2 11.7% of the variance; treatment groups also showed extensive overlap, with no clear separation of the BHB-treated group from glucose-deprived samples. Inspection of individual experimental sets (Supplementary Figure S1) confirmed that directional patterns were broadly consistent across replicate experiments despite considerable between-batch variability.

Pairwise fold-change analysis with FDR correction (Fig. 1E, F) quantified the treatment effects. In MCF-7 cells comparing BHB-supplemented to glucose-deprived conditions, 18 of 24 tested metabolites (75%) trended upward, indicating a broad rather than pathway-restricted effect of BHB. 1,4-Butanediol reached nominal significance with a 2.35-fold increase (p = 0.016), and lactic acid showed a 4.1-fold increase that did not reach nominal significance owing to high within-group variability (p = 0.11). Several additional metabolites showed nominal trends (p < 0.15), including methylamine, 2-(dimethylamino)ethanol, 2,2,2-trifluoroethane-1,1-diol, ethanolamine, and 1,3-propanediol. Glycine showed a 2.03-fold increase that did not reach significance (p = 0.25). No metabolite survived FDR correction (lowest adjusted p = 0.37), reflecting the modest sample size and high coefficient of variation (CV) typical of untargeted metabolomics. In T47D cells, the same directional analysis yielded 14 of 24 metabolites (58%) trending upward — a value essentially indistinguishable from a random 50% baseline and in stark contrast to the 75% upward trend observed in MCF-7. No metabolite reached nominal significance for the direct BHB effect (C vs B), consistent with the heatmap and PCA observations of a muted BHB response in this cell line. The metabolite with the smallest p-value, acetic acid (FC = 1.15, p = 0.15), did not approach the FDR threshold.

Two findings emerge clearly from this analysis. First, MCF-7 cells exhibit a broad metabolic response to BHB under glucose restriction—a pattern-level signal that is more informative than any individual metabolite measurement and that is consistent with BHB providing accessible metabolic carbon to these cells. Second, T47D cells do not show a comparable response, and the absence of a global metabolic shift in T47D is itself a noteworthy biological observation. Because no individual metabolite survived FDR correction in either cell line, all metabolite-specific claims that follow should be regarded as exploratory and hypothesis-generating. To clarify the apparent BHB-induced changes in selected metabolites of biological interest—particularly glycine, given its centrality in one-carbon metabolism, and the BCAA-related metabolites observed in our screen—we proceeded to targeted assays in independent cell preparations.

### 3.2 Targeted glycine and SHMT1 quantification

Given the central role of glycine in one-carbon metabolism and its apparent (though non-significant) elevation in the MCF-7 metabolomics dataset, we performed direct quantification of intracellular glycine using a fluorometric assay across a wider BHB concentration range (2.5–15 mM) in independent cell preparations. In MCF-7 cells, glycine concentrations were elevated at 5 mM BHB (51.1 ± 29.5 µM) compared with 2.5 mM BHB (27.7 ± 3.9 µM, p < 0.05), but the elevation did not persist at higher BHB concentrations and was not significantly different from the low glucose alone condition (42.0 µM; Fig. 2A). T47D cells exhibited a similar non-monotonic pattern, with glycine elevation at 5 mM BHB (41.8 ± 5.9 µM) compared with both high glucose (25.7 ± 7.8 µM, p < 0.001) and low glucose (29.6 ± 9.5 µM, p < 0.05); levels at 10 and 15 mM BHB returned to near low-glucose values (Fig. 2B). We emphasize that this profile does not constitute a dose-dependent response: increasing BHB concentration did not produce monotonically increasing glycine levels in either cell line, and elevation occurred at one intermediate concentration only.

**Figure 2.**
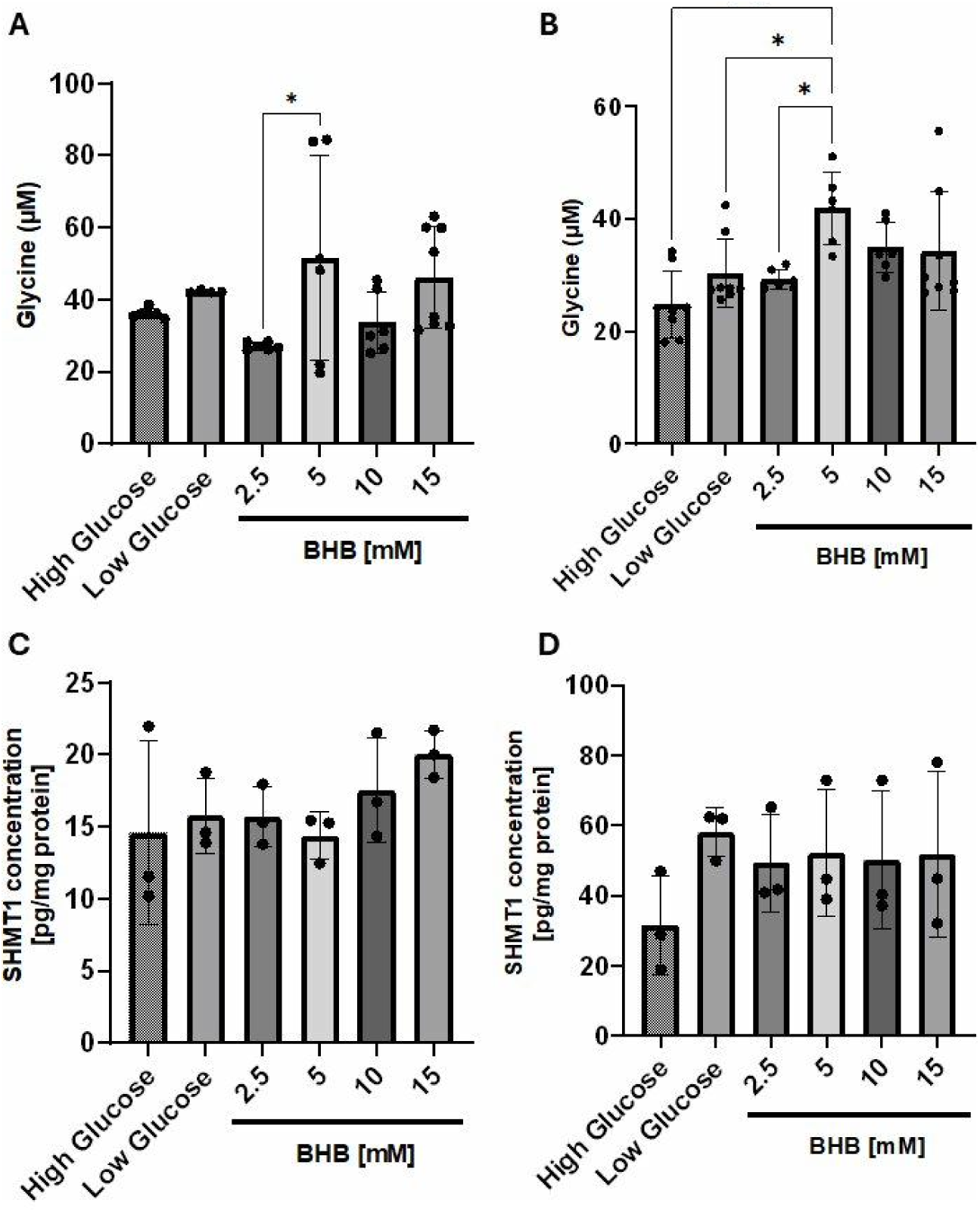
Glycine concentrations and SHMT1 protein expression in glucose-restricted breast cancer cells supplemented with β-hydroxybutyrate. (A, B) Intracellular glycine concentrations measured by fluorometric assay in MCF-7 (A) and T47D (B) cells cultured under high glucose, low glucose (5% of standard), or low glucose supplemented with 2.5, 5, 10, or 15 mM BHB for four days. (C, D) SHMT1 protein levels quantified by sandwich ELISA in MCF-7 (C) and T47D (D) cells under identical conditions. Data represent mean ± SD from three independent experiments with duplicate measurements. Statistical significance was determined by one-way ANOVA with Tukey’s post-hoc test; *p < 0.05, ***p < 0.001 for the indicated pairwise comparisons. SHMT1 protein levels did not differ significantly across treatment conditions in either cell line.

SHMT1 protein levels, quantified by ELISA in parallel cell preparations, did not differ significantly across treatment conditions in either cell line (Fig. 2C, D). MCF-7 cells displayed SHMT1 concentrations ranging from 14.4 ± 4.4 pg/mg protein (high glucose) to 17.3 ± 3.9 pg/mg protein (10 mM BHB); T47D cells showed values from 30.6 ± 11.6 pg/mg protein (high glucose) to 57.8 ± 18.3 pg/mg protein (low glucose), with overlap across all groups. The absence of change in SHMT1 protein abundance, combined with the non-monotonic glycine pattern, indicates that any glycine perturbation occurring under these conditions is not driven by transcriptional or translational regulation of SHMT1. We make no mechanistic claim about post-translational regulation of SHMT1 in this study, since we did not directly measure SHMT1 enzymatic activity or post-translational modifications.

The 10 mM BHB condition was assayed by both GC×GC-MS metabolomics (Section 3.1) and fluorometric glycine quantification (this section), allowing direct comparison. The two assays were not in quantitative agreement at this concentration. In the metabolomics analysis, MCF-7 glycine showed a non-significant 2.03-fold trend increase under BHB versus low glucose; in the targeted assay, MCF-7 glycine at 10 mM BHB (33.6 ± 13.9 µM) was numerically lower than at low glucose alone (42.0 µM). For T47D, both assays showed small upward trends with BHB that were not statistically significant. The discrepancy in MCF-7 likely reflects the different measurement principles (relative TMS-derivative peak area in metabolomics versus absolute concentration in cell lysate by fluorometric assay), the high within-group variability of the metabolomics measurement (control CV = 105%, treated CV = 62%), and the fact that the targeted assay was performed on independent biological samples rather than the same lysates used for metabolomics. We interpret this lack of concordance as evidence that glycine perturbation under these conditions, if real, is modest in magnitude relative to its baseline biological variability.

### 3.3 Total BCAA levels are elevated by BHB in T47D but not MCF-7 cells

Several BCAA-related metabolites (L-leucine, L-isoleucine, L-valine, α-hydroxyisobutyric acid) were detected in the MCF-7 metabolomics dataset, with α-hydroxyisobutyric acid additionally appearing in T47D. None reached FDR-adjusted significance in either cell line. To examine this pathway directly, we measured total intracellular BCAA concentrations across a BHB dose range (2.5–15 mM) using a colorimetric leucine dehydrogenase-based assay. We emphasize that this assay reports the sum of leucine, isoleucine, and valine and does not distinguish between the individual amino acids; resolving them would require LC-MS/MS quantification, which is a planned follow-up.

In MCF-7 cells, total BCAA concentrations remained stable across all treatment conditions, with no statistically significant differences observed between high glucose, low glucose, or any of the BHB-supplemented groups (Fig. 3A). In T47D cells, total BCAA levels were significantly elevated under 2.5 mM and 10 mM BHB conditions relative to high glucose controls (p < 0.01 for both comparisons), with values approximately doubling at peak (Fig. 3B). This pattern was not strictly dose-dependent—the 5 mM and 15 mM conditions did not reach significance—but the overall trend toward elevated BCAAs with BHB supplementation was a cell line-specific T47D phenomenon and was not observed in MCF-7.

**Figure 3.**
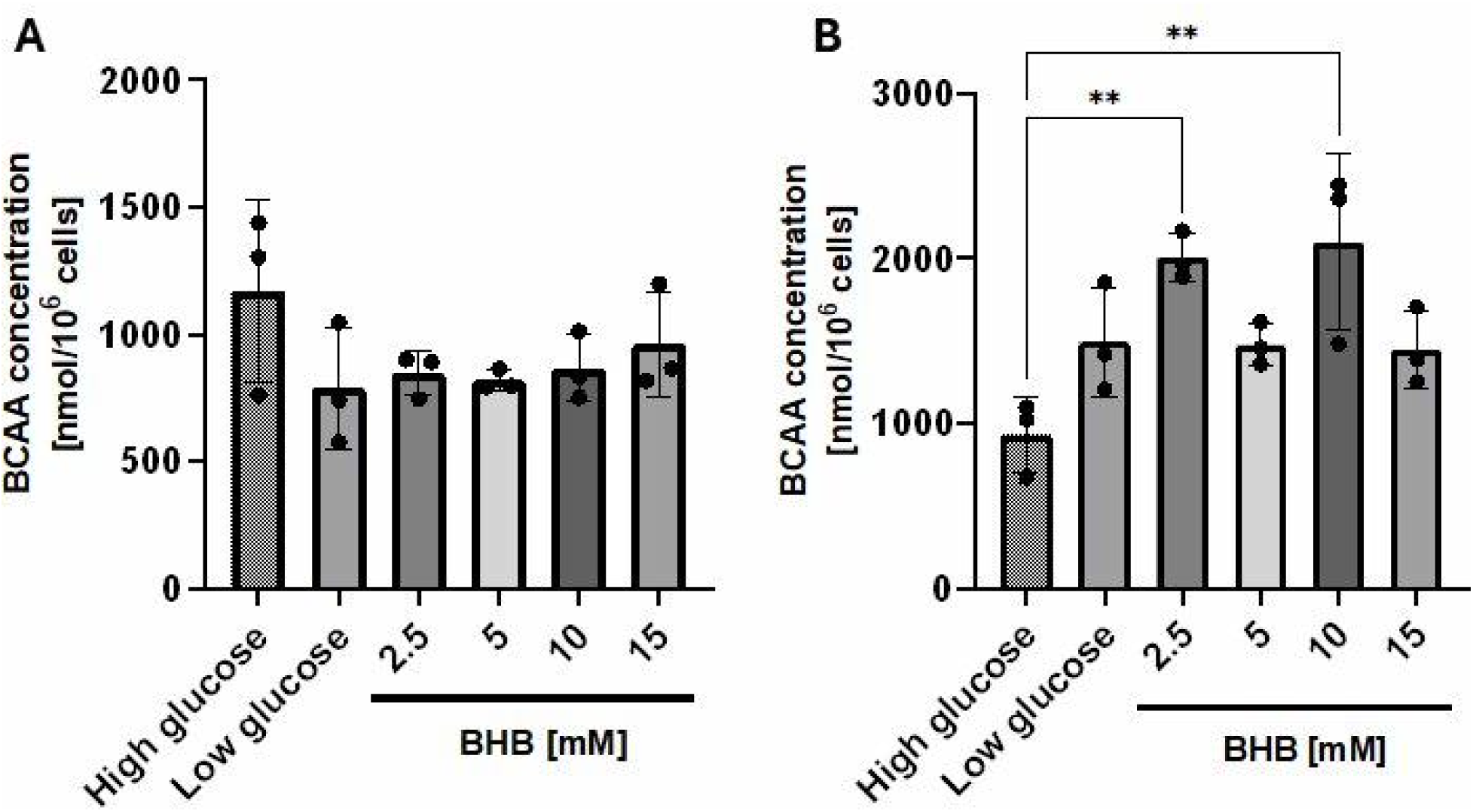
Total branched-chain amino acid (BCAA) levels in glucose-restricted breast cancer cells supplemented with β-hydroxybutyrate. Total intracellular BCAA concentrations measured by colorimetric assay in MCF-7 (A) and T47D (B) cells cultured under high glucose, low glucose (5% of standard), or low glucose with 2.5, 5, 10, or 15 mM BHB for four days. BCAA levels were quantified using a leucine dehydrogenase-based assay against an L-leucine standard curve; the assay reports the sum of leucine, isoleucine, and valine. Data represent mean ± SD from three independent experiments with triplicate measurements. Statistical significance was determined by one-way ANOVA with Tukey’s post-hoc test; **p < 0.01 versus high glucose control.

**Figure 4.**
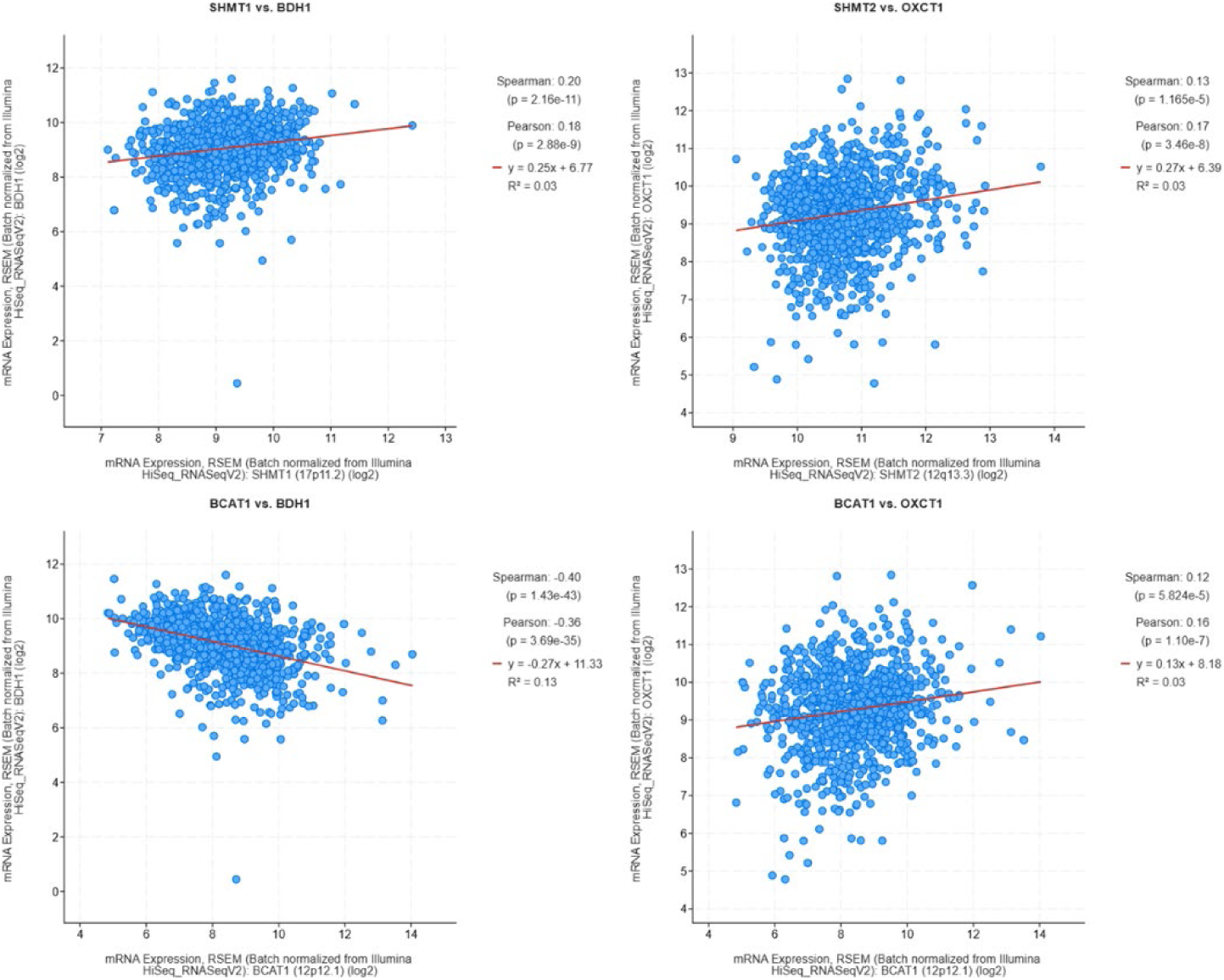
Statistically significant co-expression relationships between one-carbon/BCAA metabolism genes and ketone body catabolic genes in the TCGA Breast Invasive Carcinoma (PanCancer Atlas) cohort. Scatter plots of mRNA expression (RSEM, batch normalized, log₂) for the four gene pairs reaching both statistical significance after Bonferroni correction and a minimum correlation magnitude of r > 0.10 across 1,084 samples: SHMT1 vs. BDH1, SHMT2 vs. OXCT1, BCAT1 vs. BDH1, and BCAT1 vs. OXCT1.

This cell line-specific BCAA response is consistent with the cell line divergence observed in the global metabolomics: T47D cells, which showed little global metabolic response to BHB in the GC×GC-MS analysis, nonetheless exhibited a selective increase in total BCAAs detectable by the targeted colorimetric assay. We do not interpret this finding as a general feature of BHB action in ER+ breast cancer; rather, it appears to reflect a T47D-specific metabolic adaptation. The molecular basis of this divergence—whether due to differential expression of BCAA catabolic enzymes, transporters, or downstream utilization pathways—was not addressed in this study.

### 3.4 Selective co-expression of one-carbon and ketone body metabolism genes in breast cancer

To examine whether the metabolic relationships observed in our cell line models are reflected at the transcriptomic level in patient tumors, we performed co-expression analysis between three one-carbon and BCAA metabolism genes (SHMT1, SHMT2, BCAT1) and three ketone body catabolic genes (OXCT1, ACAT1, BDH1) in the TCGA Breast Invasive Carcinoma cohort (n = 1,084 samples; full pairwise correlation matrix in Supplementary Table S2). Most pairwise comparisons showed weak or non-significant correlations after Bonferroni correction. Four gene pairs reached both statistical significance and a minimum correlation magnitude (r > 0.10) (Fig. 4): SHMT1–BDH1 (Pearson r = 0.18, Spearman ρ = 0.20, p = 2.88 × 10⁻⁹), SHMT2–OXCT1 (r = 0.17, ρ = 0.13, p = 3.46 × 10⁻⁸), BCAT1–BDH1 (r = −0.36, ρ = −0.40, p = 3.69 × 10⁻³⁵), and BCAT1–OXCT1 (r = 0.16, ρ = 0.12, p = 1.10 × 10⁻⁷).

The most pronounced relationship was a moderate negative correlation between BCAT1 and BDH1 (Spearman ρ = −0.40, R² = 0.13), suggesting that breast tumors with high BCAA transamination capacity tend to have lower ketone body utilization, and vice versa. The remaining significant correlations were weak (R² ≈ 0.03), and we interpret them as exploratory transcriptional associations rather than evidence of direct mechanistic coupling. SHMT1 and SHMT2 did not show strong correlations with most ketone body catabolic enzymes, consistent with one-carbon and ketone body metabolism representing largely independent transcriptional programs in unstratified breast cancer.

Linear regression line in red; Pearson (r) and Spearman (ρ) correlation coefficients with p-values, and R² are annotated. The remaining five gene pairs tested did not reach the joint significance and magnitude threshold (full correlation matrix in Supplementary Table S2).

### 3.5 Coordinated downregulation of OXCT1, ACAT1, and SHMT1 in ER+ tumors compared with matched normal tissue

To examine whether the genes implicated in our metabolomics and co-expression analyses are differentially expressed in ER+ breast cancer compared with normal breast tissue, we performed paired differential expression analysis on 20 patients from the GSE58135 dataset with strictly matched ER+ primary tumor and adjacent uninvolved breast tissue samples. We analyzed nine genes involved in ketone body, one-carbon, and BCAA metabolism (Fig. 5). Three genes were significantly downregulated in tumor tissue relative to matched normal tissue: OXCT1 (log₂ fold change = −1.41, adjusted p = 5.75 × 10⁻⁷), SHMT1 (log₂ fold change = −1.31, adjusted p = 1.62 × 10⁻⁵), and ACAT1 (log₂ fold change = −1.07, adjusted p = 1.07 × 10⁻⁴). The paired analysis revealed consistent directional changes across most patients for these three genes (Fig. 5A). SHMT2, ACAT2, BDH1, BDH2, OXCT2, and BCAT1 showed no significant differential expression between tumor and normal tissue (adjusted p > 0.05; Fig. 5B).

**Figure 5.**
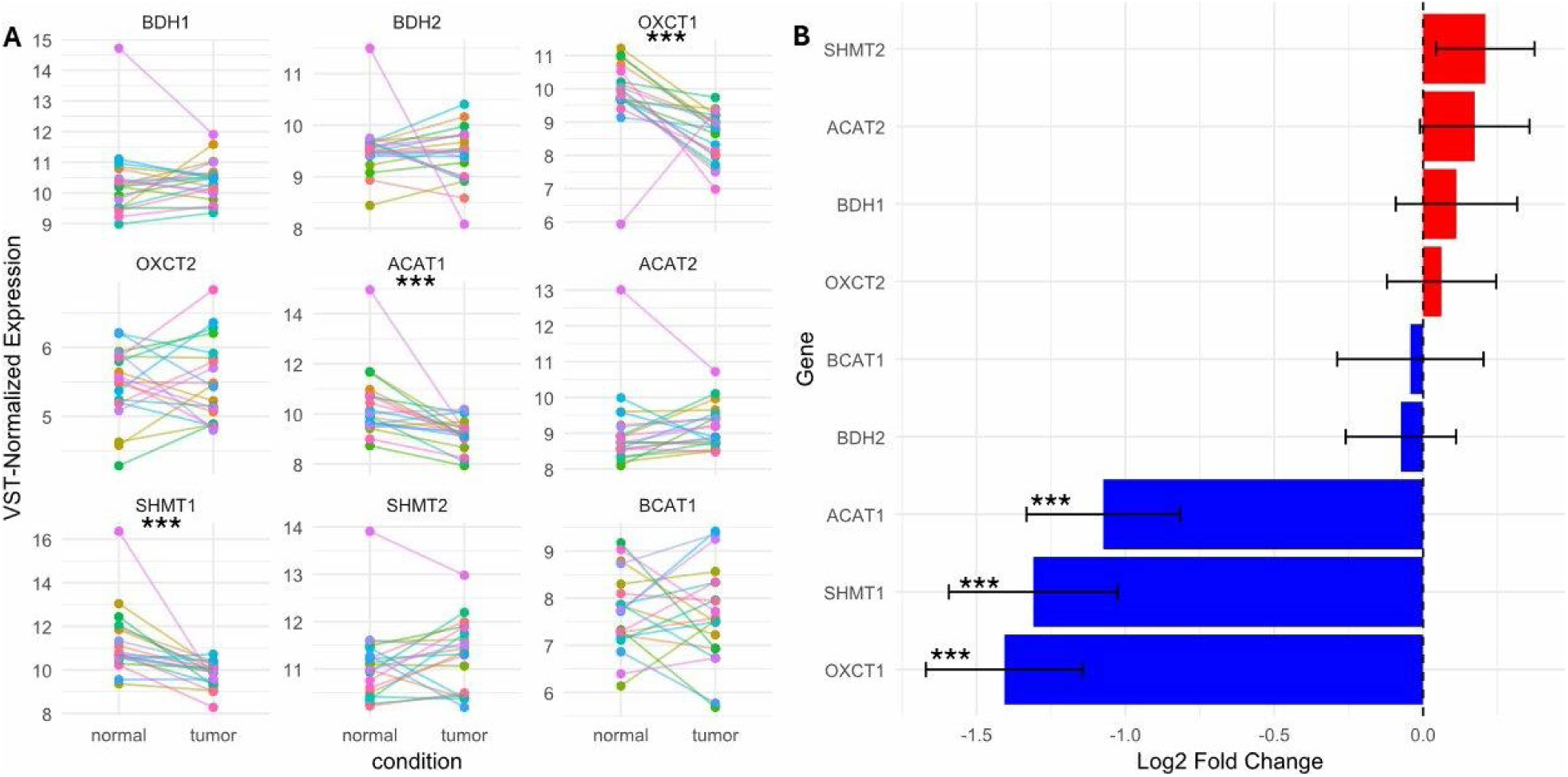
Differential expression of ketone body, one-carbon, and branched-chain amino acid metabolism genes in ER+ breast cancer tumors compared with matched adjacent normal tissue (GSE58135, n=20 paired samples). (A) Paired analysis showing VST-normalized expression levels for individual patients across normal (left) and tumor (right) tissue. Each colored line represents a single patient connecting matched normal and tumor samples; statistical significance from DESeq2 paired analysis is indicated above each panel. (B) Log₂ fold change (tumor versus normal) for each gene with standard error bars. Red bars indicate upregulation in tumors, blue bars indicate downregulation. The dashed line represents no change (log₂FC = 0). Statistical significance: *adjusted p < 0.05, **adjusted p < 0.01, ***adjusted p < 0.001 (Benjamini–Hochberg correction).

This result identifies a coordinated pattern in which ER+ tumors selectively downregulate the cytosolic one-carbon enzyme (SHMT1) and two key ketone body catabolic enzymes (OXCT1, ACAT1), while preserving expressions of their mitochondrial or alternative counterparts. The pattern is consistent across patients (Fig. 5A), in contrast to the more heterogeneous behavior seen in our in vitro experiments and suggests that reduced capacity for both cytosolic one-carbon metabolism and ketone body utilization is a recurrent feature of ER+ breast cancer biology.

## 4. Discussion

Using a combination of GC×GC-MS metabolomics, targeted biochemical assays, and patient transcriptomic analyses, we examined how two ER+ breast cancer cell lines respond to β-hydroxybutyrate supplementation under glucose restriction. The central observation of this study is that MCF-7 and T47D cells show markedly divergent metabolic responses to identical BHB treatment conditions. MCF-7 cells exhibit a broad, pattern-level metabolic shift in which the majority of detected metabolites trend upward with BHB, whereas T47D cells show essentially no comparable global response. This cell line divergence is the most reproducible feature of our dataset and was evident across metabolomics heatmaps, PCA, fold-change distributions, and the cell line-specific BCAA response in the targeted assay. We frame this divergence as the principal finding of the work and as a hypothesis-generating result regarding heterogeneity in metabolic responses to ketone bodies among ER+ breast cancer subtypes.

Heterogeneity between MCF-7 and T47D cells is well established at the proteomic and bioenergetic levels: over 160 proteins differ in abundance between the two lines [23], MCF-7 cells display higher mitochondrial reserve capacity, and T47D cells have higher glycolytic flux with greater proton leak [24]. A parsimonious interpretation of our findings is that MCF-7 cells, with their greater oxidative capacity, are better positioned to oxidize BHB-derived acetyl-CoA in the TCA cycle, leading to a broad replenishment of metabolic intermediates that would otherwise be depleted by glucose restriction. T47D cells, with their relatively higher glycolytic dependence and lower mitochondrial efficiency, may have less capacity to integrate BHB-derived carbon into their metabolic network. This interpretation is testable through measurement of ketolysis enzyme expression (BDH1, OXCT1, ACAT1) and ¹³C-BHB tracing, both of which we identify as priorities for follow-up work.

We deliberately avoid stronger mechanistic claims for two reasons. First, no individual metabolite reached FDR-adjusted significance in either cell line, and within-group coefficients of variation were high (often >50%, occasionally >100%); both observations limit confidence in any single-metabolite conclusion. Power analysis suggests that approximately 20 samples per group would be required to detect 2-fold changes at 80% power given the observed CVs, compared with the 8 to 15 samples per group available here. Second, our SHMT1 measurements show that SHMT1 protein abundance does not change across treatment conditions in either cell line, and we did not measure SHMT1 enzymatic activity, post-translational modifications, or substrate flux through one-carbon pathways. The earlier framing of these data as evidence for post-translational regulation of SHMT1 was speculative and is not supported by the present dataset; we therefore report only that any glycine perturbation observed is unaccompanied by changes in SHMT1 protein abundance, without inferring the mechanism by which glycine homeostasis might be altered.

The metabolomics screen did not identify glycine as a statistically significant treatment-responsive metabolite in either cell line after FDR correction, and the supplementary table shows glycine p-values of 0.42 (MCF-7) and 0.45 (T47D) for the three-condition comparison. The targeted fluorometric assay detected an elevation in glycine at one BHB concentration (5 mM) in both cell lines that did not persist at higher BHB concentrations and that did not represent a monotonic dose response. Furthermore, the two glycine measurements were not concordant at 10 mM BHB: metabolomics showed a non-significant upward trend in MCF-7 glycine with BHB, whereas the fluorometric assay showed numerically lower glycine at 10 mM BHB than at low glucose alone. We attribute the discrepancy to differences in measurement principle (peak area of TMS-derivative versus absolute concentration), high baseline variability in the metabolomics data, and the use of independent biological samples for the two assays. Taken together, we consider the glycine data exploratory and inconclusive with respect to a robust biological signal. Whether a small but reproducible glycine perturbation exists under BHB supplementation, and what its mechanistic basis might be, will require more sensitive and lower-variance approaches such as targeted LC-MS/MS quantification with appropriately powered sample sizes.

Total BCAA levels were elevated by BHB in T47D cells but not in MCF-7 cells. We emphasize that this finding is cell line-specific rather than general, and that our colorimetric assay reports the sum of leucine, isoleucine, and valine and cannot resolve the individual contributions of each. Although the GC×GC-MS metabolomics detected several BCAA-related features (L-leucine, L-isoleucine, L-valine, α-hydroxyisobutyric acid), none reached FDR-adjusted significance, leaving the targeted assay as the primary evidence for the T47D BCAA response. Possible explanations include reduced BCAA catabolism in T47D under metabolic stress (BCAAs serve roles in mTORC1 activation, protein synthesis, and oxidation to acetyl-CoA and succinyl-CoA [27,28]), differential BCAA transporter expression, or shifts in protein turnover; distinguishing among these will require LC-MS/MS quantification of the individual BCAAs, ¹³C-BCAA tracing, or transcriptional profiling of the BCAA catabolic pathway. Elevated BCAAs have been reported to suppress breast tumor growth and metastasis via NK cell activation [29], raising the possibility that the BCAA accumulation observed in T47D cells under BHB has functional consequences worth pursuing.

Our analysis of paired tumor-normal samples from GSE58135 identified coordinated downregulation of OXCT1, ACAT1, and SHMT1 in ER+ breast cancer compared with matched adjacent normal tissue. These changes were highly significant after FDR correction (adjusted p < 0.001 for all three genes) and were consistent across most patients in the paired analysis, distinguishing this result from the more variable cell line observations. The selective downregulation of cytosolic SHMT1 (with preserved SHMT2) is consistent with previous reports that mitochondrial SHMT2 plays the dominant role in cancer one-carbon metabolism [19,30]. The concurrent downregulation of OXCT1 and ACAT1, two enzymes central to ketone body catabolism (BHB → acetoacetate → acetyl-CoA), is particularly relevant given ongoing clinical interest in ketogenic diets as adjuvant therapy in breast cancer [31,32].

Reduced expression of OXCT1 and ACAT1 would in principle limit the capacity of ER+ tumors to oxidize ketone bodies, which could either enhance the therapeutic potential of a ketogenic diet (by creating a metabolic disadvantage for tumor cells that cannot use the alternative substrate) or limit the metabolic effect of BHB on the tumor itself. The relationship is likely complex; previous work indicated that elevated BDH1 and ACAT1 expression in breast tumors are associated with reduced overall survival [33], suggesting that tumors retaining ketone body utilization capacity may behave more aggressively. We propose that tumor-level expression of OXCT1, ACAT1, and SHMT1 deserves further evaluation as a candidate biomarker panel for stratifying patients who may benefit from ketogenic dietary interventions—a question that will require prospective clinical study in a larger and more heterogeneous patient population.

In our TCGA co-expression analysis, we restricted reporting to the four gene pairs reaching both Bonferroni-corrected significance and a minimum correlation magnitude. The strongest finding—a moderate negative correlation between BCAT1 and BDH1 (Spearman ρ = −0.40)—suggests an inverse relationship between BCAA transamination and ketone body utilization at the transcriptional level. The remaining significant correlations were weak (R² ≈ 0.03) and may reflect broad transcriptional co-regulation programs rather than direct functional coupling; we therefore frame them as exploratory and as candidate hypotheses for follow-up rather than as direct support for any specific mechanism.

Several limitations of this study should be acknowledged. First, the GC×GC-MS metabolomics analysis showed substantial inter-experimental batch variability and high within-group coefficients of variation, limiting our ability to detect individual treatment-responsive metabolites after FDR correction. We addressed this by focusing on pattern-level effects (proportion of metabolites trending upward) rather than individual significance, and by performing follow-up targeted assays in independent biological samples; nonetheless, the metabolomics findings should be regarded as exploratory. A larger sample size (n ≈ 20 per group) and improved batch correction would substantially strengthen future analyses. Second, our study was limited to two ER+ breast cancer cell lines cultured in monolayer; the cell line divergence we observed underscores the need to extend these analyses to additional ER+ models, to triple-negative or HER2-amplified subtypes for comparison, and ultimately to three-dimensional culture systems and patient-derived samples. Third, the BHB concentrations used in our targeted assays (2.5–15 mM) include values that span physiological (∼0.5–1 mM mild ketosis), therapeutic ketogenic diet (∼2–5 mM), and supraphysiological ranges; the non-monotonic patterns observed at intermediate concentrations make it important not to extrapolate uncritically from in vitro doses to clinical conditions. Fourth, the colorimetric BCAA assay does not distinguish leucine, isoleucine, and valine, and we did not perform direct enzymatic activity, post-translational modification, or flux measurements on SHMT1. Finally, while the GSE58135 paired analysis is statistically robust within its sample size, the cohort is heterogeneous in clinical and molecular characteristics; stratification by molecular subtype, stage, or treatment history might reveal stronger or more specific associations.

## 5. Conclusions

This study reframes a previous observation of broad metabolic effects of β-hydroxybutyrate supplementation under glucose restriction in ER+ breast cancer cells around the central feature most strongly supported by the data: cell line heterogeneity. MCF-7 and T47D cells, both ER+ breast cancer models in widespread experimental use, respond divergently to identical BHB treatment conditions, with MCF-7 showing a broad pattern-level metabolic shift and T47D showing essentially no global response. Individual metabolite-specific claims, including those previously made about glycine and one-carbon metabolism, do not survive FDR correction in our metabolomics data and should be regarded as exploratory. Total BCAA elevation under BHB is a T47D-specific phenomenon. In paired patient samples, OXCT1, ACAT1, and SHMT1 are coordinately downregulated in ER+ tumors relative to matched normal tissue, suggesting these enzymes may be relevant for stratifying patients in clinical investigations of ketogenic interventions. Together, our findings argue for caution in generalizing in vitro metabolic responses to BHB across ER+ cell line models and identify candidate transcriptional markers for further translational evaluation. Future work should include direct measurement of ketolysis enzyme activity in MCF-7 versus T47D, ¹³C-BHB and ¹³C-BCAA tracing to define carbon fates in each cell line, individual BCAA quantification by LC-MS/MS, and prospective evaluation of OXCT1/ACAT1/SHMT1 expression as candidate biomarkers in ketogenic diet clinical trials.

## Supplementary information

**Supplementary Table S1.** GC×GC-MS metabolite quantification in MCF-7 and T47D breast cancer cells. Pairwise Welch’s t-test results with Benjamini–Hochberg FDR correction for each of three pairwise comparisons (low glucose vs. high glucose; BHB-supplemented vs. high glucose; BHB-supplemented vs. low glucose) in each cell line. Reports log₂ fold change, fold change, raw p-value, adjusted p-value, within-group coefficients of variation, and direction of change for each detected metabolite.

**Supplementary Table S2.** Complete pairwise correlation matrix for SHMT1, SHMT2, and BCAT1 versus OXCT1, ACAT1, and BDH1 in the TCGA Breast Invasive Carcinoma cohort (n = 1,084 samples). Pearson and Spearman coefficients with corresponding p-values are reported for all nine gene pairs; the four pairs meeting both Bonferroni-corrected significance (α = 0.0056) and a minimum correlation magnitude (r > 0.10) are highlighted and shown individually in Figure 4 of the main manuscript.

**Supplementary Figure S1.** Individual experimental set PCA analysis showing PCA scores (left panels per set) and loadings (right panels per set) for MCF-7 (sets 3, 4, 5) and T47D (sets 3, 4, 5) breast cancer cells under high glucose, low glucose, and low glucose with BHB conditions. Treatment-dependent directional separation is consistent across individual experimental sets despite between-batch technical variability.

## Declarations

### Ethics approval and consent to participate

This study did not involve human subjects or animal models. All experiments were conducted using commercially available cell lines or publicly available data. Institutional Review Board (IRB) and Institutional Animal Care and Use Committee (IACUC) approval were not required.

### Consent for publication

Not applicable.

### Availability of data and materials

The raw data of the findings of this study are available from the corresponding author upon reasonable request. The TCGA data analyzed are available in the cBioPortal repository (https://www.cbioportal.org). The GSE58135 dataset is available from the Gene Expression Omnibus (https://www.ncbi.nlm.nih.gov/geo/query/acc.cgi?acc=GSE58135).

### Competing interests

The authors declare no potential conflicts of interest.

## Funding

This project was supported by grants from the National Institutes of Health (NIH), National Institute of General Medical Sciences (NIGMS), IDeA Networks of Biomedical Research Excellence (INBRE), Award number P20GM103466. The content is solely the responsibility of the authors and does not necessarily represent the official views of the National Institutes of Health.

## Authors’ contributions

CC and NG contributed equally to this work. CC performed metabolomics experiments and data analysis. NG supervised undergraduate students, performed data analysis, and created figures. RM, SB, TS, and LT performed cell culture and biochemical assays. KAPU supervised and provided technical expertise for the GC×GC-MS experimental setup and data analysis, and edited the manuscript. MW conceived the study, performed RNA-seq analysis and bioinformatics, supervised the research, and wrote the manuscript. All authors read and approved the final manuscript.

## Acknowledgements

Not applicable.

## Notes

https://www.cbioportal.org

https://www.ncbi.nlm.nih.gov/geo/query/acc.cgi?acc=GSE58135

### Competing Interest Statement

The authors have declared no competing interest.

