## Supplementary Table S1 for "β-Hydroxybutyrate elicits divergent metabolic responses between MCF-7 and T47D ER+ breast cancer cells under glucose restriction"

### Supplementary Table S1. GC×GC-MS metabolite quantification in MCF-7 and T47D breast cancer cells

*Welch's t-tests with Benjamini–Hochberg FDR correction. MCF-7 analysis (n=8 per group); T47D analysis (n=14–15 per group). Significance coding:* ***red*** *= FDR p<0.05 with |FC|>1.5;* ***pink*** *= nominal p<0.05;* ***yellow*** *= trend (p<0.15 with |FC|>1.5). FC = fold change; CV = coefficient of variation.*

#### A. MCF-7 cells (n=8 per group)

##### A1. Low Glucose vs High Glucose (B vs A) — 46 metabolites tested

| Metabolite | $Log_2FC$ | FC | p-value | Adj. p-value | CV Ctrl (%) | CV Trt (%) | Direction |
|---|---|---|---|---|---|---|---|
| Isopropyl alcohol | 0.85 | 1.80 | 0.1634 | 0.9434 | 91.1 | 68.7 | ↑ |
| p-Xylene | 0.32 | 1.25 | 0.1901 | 0.9434 | 24.8 | 34.7 | ↑ |
| Pentane, 3-methyl- | 0.53 | 1.44 | 0.2006 | 0.9434 | 3.2 | 28.4 | ↑ |
| n-Hexane | 0.13 | 1.10 | 0.2034 | 0.9434 | 1.0 | 8.1 | ↑ |
| Cyclopentane, methyl- | 0.24 | 1.18 | 0.2132 | 0.9434 | 4.9 | 15.2 | ↑ |
| Pentane, 2-methyl- | -0.10 | 0.93 | 0.2541 | 0.9434 | 7.3 | 4.9 | ↓ |
| Lactic Acid | -1.44 | 0.37 | 0.2574 | 0.9434 | 141.7 | 101.1 | ↓ |
| Pentane, 2-methyl- | 0.21 | 1.16 | 0.2624 | 0.9434 | 16.4 | 29.0 | ↑ |
| 2-(Dimethylamino)ethanol | 0.79 | 1.73 | 0.3100 | 0.9434 | 47.1 | 53.5 | ↑ |
| 1,4-Butanediol | -0.72 | 0.61 | 0.3195 | 0.9434 | 69.6 | 90.1 | ↓ |
| Pyrazine | 0.32 | 1.25 | 0.3463 | 0.9434 | 55.4 | 34.6 | ↑ |
| L-Valine | -0.92 | 0.53 | 0.3771 | 0.9434 | 72.2 | 46.5 | ↓ |
| Silanol, trimethyl- | 0.90 | 1.86 | 0.3838 | 0.9434 | 33.4 | 72.2 | ↑ |
| Isopropyl alcohol | -0.44 | 0.74 | 0.4005 | 0.9434 | 28.4 | 52.9 | ↓ |
| Acetic acid | -1.45 | 0.37 | 0.4106 | 0.9434 | 122.2 | 235.5 | ↓ |
| 1,3-Propanediol | 0.99 | 1.99 | 0.4116 | 0.9434 | 33.8 | 83.5 | ↑ |
| Butanoic acid, 4-hydroxy- | 1.14 | 2.21 | 0.4302 | 0.9434 | 53.5 | 96.1 | ↑ |
| Pyrazine | 0.18 | 1.13 | 0.4565 | 0.9434 | 8.1 | 21.5 | ↑ |
| Lactic Acid | 0.40 | 1.32 | 0.4839 | 0.9434 | 59.9 | 29.4 | ↑ |
| Ethylbenzene | 0.32 | 1.24 | 0.5066 | 0.9434 | 36.1 | 36.3 | ↑ |
| Ethanolamine | -0.64 | 0.64 | 0.5311 | 0.9434 | 97.5 | 79.4 | ↓ |
| 2,2,2-Trifluoroethane-1,1-diol, 2TMS | -0.96 | 0.51 | 0.5421 | 0.9434 | 166.2 | 234.3 | ↓ |
| Glycine | -0.64 | 0.64 | 0.5925 | 0.9434 | 89.3 | 117.4 | ↓ |
| 2-(Dimethylamino)ethanol | -0.49 | 0.71 | 0.5993 | 0.9434 | 134.4 | 89.6 | ↓ |
| Methoxyamine | -0.23 | 0.86 | 0.6189 | 0.9434 | 36.3 | 33.8 | ↓ |
| Butanoic acid, 4-hydroxy- | 0.73 | 1.66 | 0.6204 | 0.9434 | 204.7 | 178.4 | ↑ |
| Cyclopentane, methyl- | 0.09 | 1.07 | 0.6446 | 0.9434 | 29.9 | 23.8 | ↑ |
| Ethylbenzene | -0.50 | 0.71 | 0.6802 | 0.9434 | 158.3 | 132.1 | ↓ |
| Pentane, 3-methyl- | 0.07 | 1.05 | 0.6903 | 0.9434 | 21.0 | 26.3 | ↑ |
| Methylamine, N,N-dimethyl- | -0.24 | 0.85 | 0.6903 | 0.9434 | 57.7 | 1.0 | ↓ |
| Ethylene glycol | 0.09 | 1.06 | 0.7072 | 0.9434 | 6.1 | 22.6 | ↑ |
| Pyridine | -0.23 | 0.85 | 0.7602 | 0.9434 | 93.4 | 72.0 | ↓ |
| Methylamine, N,N-dimethyl- | -0.32 | 0.80 | 0.7646 | 0.9434 | 71.3 | 108.0 | ↓ |
| Silanol, trimethyl- | 0.26 | 1.20 | 0.7693 | 0.9434 | 131.8 | 100.4 | ↑ |
| 1,4-Butanediol | -0.08 | 0.95 | 0.7858 | 0.9434 | 22.6 | 22.7 | ↓ |
| Methoxyamine | -0.36 | 0.78 | 0.7914 | 0.9434 | 137.1 | 186.6 | ↓ |
| L-Alanine | -0.22 | 0.86 | 0.8008 | 0.9434 | 72.6 | 59.9 | ↓ |
| Pyridine | -0.08 | 0.95 | 0.8254 | 0.9434 | 15.2 | 36.1 | ↓ |

| Metabolite | $Log_2FC$ | FC | p-value | Adj. p-value | CV Ctrl (%) | CV Trt (%) | Direction |
|---|---|---|---|---|---|---|---|
| Acetic acid | 0.09 | 1.07 | 0.8495 | 0.9434 | 52.9 | 11.4 | ↑ |
| Methylamine | -0.08 | 0.94 | 0.8556 | 0.9434 | 21.5 | 47.1 | ↓ |
| Ethylene glycol | 0.11 | 1.08 | 0.8558 | 0.9434 | 86.3 | 63.5 | ↑ |
| 2,2,2-Trifluoroethane-1,1-diol, 2TMS | -0.11 | 0.93 | 0.8613 | 0.9434 | 63.2 | 6.1 | ↓ |
| 1,3-Propanediol | -0.08 | 0.94 | 0.9393 | 0.9676 | 154.3 | 136.1 | ↓ |
| Methylamine | -0.03 | 0.98 | 0.9434 | 0.9676 | 44.3 | 46.8 | ↓ |
| p-Xylene | -0.04 | 0.97 | 0.9553 | 0.9676 | 11.4 | 72.6 | ↓ |
| n-Hexane | 0.00 | 1.00 | 0.9676 | 0.9676 | 7.4 | 10.0 | ↑ |

#### A2. BHB + Low Glucose vs High Glucose (C vs A) — 23 metabolites tested

| Metabolite | $Log_2FC$ | FC | p-value | Adj. p-value | CV Ctrl (%) | CV Trt (%) | Direction |
|---|---|---|---|---|---|---|---|
| Methylamine | 0.60 | 1.52 | 0.0776 | 0.7061 | 44.3 | 23.8 | ↑ |
| Silanol, trimethyl- | 0.97 | 1.95 | 0.2087 | 0.7061 | 131.8 | 69.9 | ↑ |
| Pyrazine | 0.43 | 1.35 | 0.2192 | 0.7061 | 55.4 | 35.9 | ↑ |
| p-Xylene | 0.26 | 1.20 | 0.2209 | 0.7061 | 24.8 | 30.2 | ↑ |
| Butanoic acid, 4-hydroxy- | 1.65 | 3.15 | 0.2723 | 0.7061 | 204.7 | 139.3 | ↑ |
| 1,4-Butanediol | 0.51 | 1.43 | 0.2881 | 0.7061 | 69.6 | 31.9 | ↑ |
| 1,3-Propanediol | 0.94 | 1.92 | 0.2938 | 0.7061 | 154.3 | 87.7 | ↑ |
| 2-(Dimethylamino)ethanol | 0.83 | 1.77 | 0.3147 | 0.7061 | 134.4 | 90.6 | ↑ |
| Isopropyl alcohol | 0.86 | 1.81 | 0.3303 | 0.7061 | 91.1 | 104.8 | ↑ |
| Methylamine, N,N-dimethyl- | 0.62 | 1.54 | 0.3353 | 0.7061 | 71.3 | 55.3 | ↑ |
| Pentane, 2-methyl- | 0.14 | 1.10 | 0.3377 | 0.7061 | 16.4 | 20.8 | ↑ |
| Methoxyamine | 0.73 | 1.66 | 0.4426 | 0.7443 | 137.1 | 79.2 | ↑ |
| 2,2,2-Trifluoroethane-1,1-diol, 2TMS | 0.77 | 1.70 | 0.4451 | 0.7443 | 166.2 | 104.7 | ↑ |
| Ethylene glycol | 0.40 | 1.32 | 0.4530 | 0.7443 | 86.3 | 45.0 | ↑ |
| Lactic Acid | 0.60 | 1.52 | 0.5264 | 0.7675 | 141.7 | 115.9 | ↑ |
| Ethanolamine | 0.44 | 1.36 | 0.5919 | 0.7675 | 97.5 | 67.4 | ↑ |
| Ethylbenzene | 0.48 | 1.40 | 0.6347 | 0.7675 | 158.3 | 111.4 | ↑ |
| n-Hexane | -0.02 | 0.98 | 0.6506 | 0.7675 | 7.4 | 6.9 | ↓ |
| Pyridine | -0.38 | 0.77 | 0.6528 | 0.7675 | 93.4 | 92.0 | ↓ |
| Glycine | 0.38 | 1.30 | 0.6674 | 0.7675 | 89.3 | 69.5 | ↑ |
| Cyclopentane, methyl- | -0.05 | 0.97 | 0.8478 | 0.9285 | 29.9 | 36.8 | ↓ |
| Pentane, 3-methyl- | -0.02 | 0.99 | 0.9267 | 0.9596 | 21.0 | 26.7 | ↓ |
| Acetic acid | -0.06 | 0.96 | 0.9596 | 0.9596 | 122.2 | 119.0 | ↓ |

#### A3. BHB + Low Glucose vs Low Glucose (C vs B) — 24 metabolites tested [KEY COMPARISON, COMPLETE]

| Metabolite | $Log_2FC$ | FC | p-value | Adj. p-value | CV Ctrl (%) | CV Trt (%) | Direction |
|---|---|---|---|---|---|---|---|
| 1,4-Butanediol | 1.23 | 2.35 | 0.0155 | 0.3726 | 90.1 | 31.9 | ↑ |
| Methylamine | 0.63 | 1.55 | 0.0568 | 0.5723 | 46.8 | 23.8 | ↑ |
| Lactic Acid | 2.05 | 4.13 | 0.1098 | 0.5723 | 101.1 | 115.9 | ↑ |
| 2-(Dimethylamino)ethanol | 1.31 | 2.48 | 0.1167 | 0.5723 | 89.6 | 90.6 | ↑ |
| 2,2,2-Trifluoroethane-1,1-diol, 2TMS | 1.73 | 3.32 | 0.1511 | 0.5723 | 234.3 | 104.7 | ↑ |
| Ethanolamine | 1.09 | 2.12 | 0.1655 | 0.5723 | 79.4 | 67.4 | ↑ |

| Metabolite | $Log_2FC$ | FC | p-value | Adj. p-value | CV Ctrl (%) | CV Trt (%) | Direction |
|---|---|---|---|---|---|---|---|
| 1,3-Propanediol | 1.02 | 2.03 | 0.2384 | 0.5723 | 136.1 | 87.7 | ↑ |
| Glycine | 1.02 | 2.03 | 0.2452 | 0.5723 | 117.4 | 69.5 | ↑ |
| Methoxyamine | 1.08 | 2.12 | 0.2634 | 0.5723 | 186.6 | 79.2 | ↑ |
| Silanol, trimethyl- | 0.71 | 1.63 | 0.2795 | 0.5723 | 100.4 | 69.9 | ↑ |
| Methylamine, N,N-dimethyl- | 0.94 | 1.92 | 0.3023 | 0.5723 | 108.0 | 55.3 | ↑ |
| Ethylbenzene | 0.98 | 1.97 | 0.3069 | 0.5723 | 132.1 | 111.4 | ↑ |
| Acetic acid | 1.39 | 2.62 | 0.3100 | 0.5723 | 235.5 | 119.0 | ↑ |
| Butanoic acid, 4-hydroxy- | 0.92 | 1.89 | 0.4657 | 0.7983 | 178.4 | 139.3 | ↑ |
| Ethylene glycol | 0.29 | 1.22 | 0.5134 | 0.8011 | 63.5 | 45.0 | ↑ |
| Cyclopentane, methyl- | -0.14 | 0.91 | 0.5390 | 0.8011 | 23.8 | 36.8 | ↓ |
| L-Valine | 0.36 | 1.28 | 0.5723 | 0.8011 | 43.2 | 70.5 | ↑ |
| Pentane, 3-methyl- | -0.09 | 0.94 | 0.6576 | 0.8011 | 26.3 | 26.7 | ↓ |
| n-Hexane | -0.03 | 0.98 | 0.6773 | 0.8011 | 10.0 | 6.9 | ↓ |
| Pentane, 2-methyl- | -0.07 | 0.95 | 0.6939 | 0.8011 | 29.0 | 20.8 | ↓ |
| Pyrazine | 0.11 | 1.08 | 0.7010 | 0.8011 | 34.6 | 35.9 | ↑ |
| Pyridine | -0.14 | 0.91 | 0.8152 | 0.8569 | 72.0 | 92.0 | ↓ |
| p-Xylene | -0.05 | 0.96 | 0.8212 | 0.8569 | 34.7 | 30.2 | ↓ |
| Isopropyl alcohol | 0.01 | 1.01 | 0.9906 | 0.9906 | 68.7 | 104.8 | ↑ |

### B. T47D cells (all sets, n=14–15 per group)

#### B1. Low Glucose vs High Glucose (B vs A) — 24 metabolites tested

| Metabolite | $Log_2FC$ | FC | p-value | Adj. p-value | CV Ctrl (%) | CV Trt (%) | Direction |
|---|---|---|---|---|---|---|---|
| Nonadecane | 0.19 | 1.14 | 0.0105 | 0.2519 | 13.3 | 12.5 | ↑ |
| Propanedioic acid | 0.42 | 1.34 | 0.0342 | 0.2749 | 24.9 | 38.8 | ↑ |
| Heptacosane | 0.15 | 1.11 | 0.0344 | 0.2749 | 13.1 | 12.1 | ↑ |
| Nonadecane | 0.12 | 1.09 | 0.0894 | 0.4755 | 13.1 | 12.4 | ↑ |
| Pentane, 2-methyl- | 0.07 | 1.05 | 0.1058 | 0.4755 | 4.9 | 8.9 | ↑ |
| 2,4,4,6-Tetramethyl-[1,3,2]dioxaborinane | -2.02 | 0.25 | 0.1189 | 0.4755 | 165.8 | 150.8 | ↓ |
| Toluene | -3.36 | 0.10 | 0.1607 | 0.5044 | 227.0 | 66.4 | ↓ |
| 2,4-Di-tert-butylphenol | 0.10 | 1.07 | 0.1730 | 0.5044 | 15.0 | 12.5 | ↑ |
| Hydroxylamine, O-methyl- | 0.10 | 1.07 | 0.1984 | 0.5044 | 11.6 | 15.0 | ↑ |
| Cyclopentane, methyl- | -3.02 | 0.12 | 0.2102 | 0.5044 | 248.6 | 125.5 | ↓ |
| 1,1-Dimethylethanol | -0.44 | 0.74 | 0.2749 | 0.5997 | 81.4 | 44.9 | ↓ |
| n-Hexane | 0.19 | 1.14 | 0.3353 | 0.6706 | 37.7 | 33.1 | ↑ |
| Acetic acid | -0.11 | 0.92 | 0.4187 | 0.6910 | 29.6 | 19.7 | ↓ |
| β-D-(-)-Ribopyranose | 0.06 | 1.04 | 0.4346 | 0.6910 | 13.5 | 14.1 | ↑ |
| Glycine | -0.50 | 0.71 | 0.4365 | 0.6910 | 126.6 | 79.7 | ↓ |
| Methylphosphonic acid | 0.26 | 1.20 | 0.4728 | 0.6910 | 59.7 | 70.5 | ↑ |
| α-Hydroxyisobutyric acid | 0.08 | 1.06 | 0.4895 | 0.6910 | 19.8 | 21.4 | ↑ |
| Butanoic acid, 2-methylbutyl ester | 0.07 | 1.05 | 0.5328 | 0.7104 | 20.3 | 21.6 | ↑ |
| N,N-Dimethyltrifluoroacetamide | 0.08 | 1.06 | 0.5845 | 0.7384 | 23.7 | 28.5 | ↑ |
| Hexane, 2-chloro- | -1.00 | 0.50 | 0.6364 | 0.7637 | 353.2 | 335.0 | ↓ |
| Glycerol | 0.05 | 1.03 | 0.6724 | 0.7685 | 21.2 | 21.2 | ↑ |
| Lactic Acid | -0.12 | 0.92 | 0.7527 | 0.8211 | 84.5 | 53.3 | ↓ |
| Acetaldehyde | -0.02 | 0.99 | 0.7928 | 0.8272 | 14.7 | 12.1 | ↓ |
| Glycine | 0.01 | 1.01 | 0.9000 | 0.9000 | 19.3 | 15.6 | ↑ |

#### B2. BHB + Low Glucose vs High Glucose (C vs A) — 24 metabolites tested

| Metabolite | $Log_2FC$ | FC | p-value | Adj. p-value | CV Ctrl (%) | CV Trt (%) | Direction |
|---|---|---|---|---|---|---|---|
| α-Hydroxyisobutyric acid | 0.27 | 1.20 | 0.0606 | 0.4219 | 19.8 | 27.3 | ↑ |
| β-D-(-)-Ribopyranose | 0.16 | 1.12 | 0.1020 | 0.4219 | 13.5 | 19.3 | ↑ |
| Butanoic acid, 2-methylbutyl ester | 0.22 | 1.17 | 0.1046 | 0.4219 | 20.3 | 26.5 | ↑ |
| N,N-Dimethyltrifluoroacetamide | 0.27 | 1.20 | 0.1099 | 0.4219 | 23.7 | 32.1 | ↑ |
| Propanedioic acid | 0.30 | 1.23 | 0.1216 | 0.4219 | 24.9 | 38.3 | ↑ |
| Methylphosphonic acid | 0.77 | 1.70 | 0.1646 | 0.4219 | 59.7 | 100.2 | ↑ |
| n-Hexane | 0.26 | 1.20 | 0.1831 | 0.4219 | 37.7 | 32.7 | ↑ |
| Heptacosane | 0.10 | 1.07 | 0.2073 | 0.4219 | 13.1 | 15.5 | ↑ |
| Nonadecane | 0.10 | 1.07 | 0.2082 | 0.4219 | 13.3 | 15.5 | ↑ |
| Nonadecane | 0.10 | 1.07 | 0.2093 | 0.4219 | 13.1 | 15.5 | ↑ |
| 1,1-Dimethylethanol | -0.50 | 0.71 | 0.2153 | 0.4219 | 81.4 | 31.6 | ↓ |
| Cyclopentane, methyl- | -2.91 | 0.13 | 0.2155 | 0.4219 | 248.6 | 133.4 | ↓ |
| Toluene | -2.20 | 0.22 | 0.2285 | 0.4219 | 227.0 | 235.3 | ↓ |
| 2,4-Di-tert-butylphenol | 0.10 | 1.07 | 0.2527 | 0.4332 | 15.0 | 15.9 | ↑ |

| Metabolite | $Log_2FC$ | FC | p-value | Adj. p-value | CV Ctrl (%) | CV Trt (%) | Direction |
|---|---|---|---|---|---|---|---|
| Hexane, 2-chloro- | 2.59 | 6.02 | 0.2850 | 0.4560 | 353.2 | 274.6 | ↑ |
| Hydroxylamine, O-methyl- | 0.05 | 1.04 | 0.4269 | 0.6233 | 11.6 | 12.9 | ↑ |
| Glycerol | 0.10 | 1.07 | 0.4415 | 0.6233 | 21.2 | 25.0 | ↑ |
| 2,4,4,6-Tetramethyl-[1,3,2]dioxaborinane | -0.78 | 0.58 | 0.5171 | 0.6895 | 165.8 | 295.7 | ↓ |
| Lactic Acid | -0.22 | 0.86 | 0.5898 | 0.7124 | 84.5 | 56.6 | ↓ |
| Acetic acid | 0.08 | 1.06 | 0.5937 | 0.7124 | 29.6 | 28.1 | ↑ |
| Pentane, 2-methyl- | 0.01 | 1.01 | 0.6751 | 0.7716 | 4.9 | 7.4 | ↑ |
| Glycine | 0.11 | 1.08 | 0.8768 | 0.9433 | 126.6 | 133.5 | ↑ |
| Acetaldehyde | 0.01 | 1.01 | 0.9274 | 0.9433 | 14.7 | 14.6 | ↑ |
| Glycine | 0.01 | 1.01 | 0.9433 | 0.9433 | 19.3 | 18.4 | ↑ |

#### B3. BHB + Low Glucose vs Low Glucose (C vs B) — 24 metabolites tested [KEY COMPARISON]

| Metabolite | $Log_2FC$ | FC | p-value | Adj. p-value | CV Ctrl (%) | CV Trt (%) | Direction |
|---|---|---|---|---|---|---|---|
| Acetic acid | 0.20 | 1.15 | 0.1546 | 0.8687 | 19.7 | 28.1 | ↑ |
| α-Hydroxyisobutyric acid | 0.19 | 1.14 | 0.1739 | 0.8687 | 21.4 | 27.3 | ↑ |
| Hexane, 2-chloro- | 3.59 | 12.03 | 0.2353 | 0.8687 | 335.0 | 274.6 | ↑ |
| Nonadecane | -0.09 | 0.94 | 0.2553 | 0.8687 | 12.5 | 15.5 | ↓ |
| Pentane, 2-methyl- | -0.05 | 0.97 | 0.2576 | 0.8687 | 8.9 | 7.4 | ↓ |
| Butanoic acid, 2-methylbutyl ester | 0.15 | 1.11 | 0.2608 | 0.8687 | 21.6 | 26.5 | ↑ |
| N,N-Dimethyltrifluoroacetamide | 0.19 | 1.14 | 0.2687 | 0.8687 | 28.5 | 32.1 | ↑ |
| β-D-(-)-Ribopyranose | 0.10 | 1.07 | 0.2906 | 0.8687 | 14.1 | 19.3 | ↑ |
| Methylphosphonic acid | 0.51 | 1.42 | 0.3289 | 0.8687 | 70.5 | 100.2 | ↑ |
| Glycine | 0.61 | 1.53 | 0.3777 | 0.8687 | 79.7 | 133.5 | ↑ |
| Toluene | 1.16 | 2.24 | 0.3981 | 0.8687 | 66.4 | 235.3 | ↑ |
| 2,4,4,6-Tetramethyl-[1,3,2]dioxaborinane | 1.23 | 2.35 | 0.4886 | 0.9239 | 150.8 | 295.7 | ↑ |
| Heptacosane | -0.05 | 0.97 | 0.5252 | 0.9239 | 12.1 | 15.5 | ↓ |
| Propanedioic acid | -0.12 | 0.92 | 0.5560 | 0.9239 | 38.8 | 38.3 | ↓ |
| Hydroxylamine, O-methyl- | -0.04 | 0.97 | 0.5898 | 0.9239 | 15.0 | 12.9 | ↓ |
| n-Hexane | 0.08 | 1.05 | 0.6701 | 0.9239 | 33.1 | 32.7 | ↑ |
| Glycerol | 0.05 | 1.04 | 0.6872 | 0.9239 | 21.2 | 25.0 | ↑ |
| Acetaldehyde | 0.03 | 1.02 | 0.7162 | 0.9239 | 12.1 | 14.6 | ↑ |
| Lactic Acid | -0.10 | 0.93 | 0.7430 | 0.9239 | 53.3 | 56.6 | ↓ |
| 1,1-Dimethylethanol | -0.06 | 0.96 | 0.7887 | 0.9239 | 44.9 | 31.6 | ↓ |
| Nonadecane | -0.02 | 0.99 | 0.8084 | 0.9239 | 12.4 | 15.5 | ↓ |
| Cyclopentane, methyl- | 0.12 | 1.08 | 0.8703 | 0.9494 | 125.5 | 133.4 | ↑ |
| 2,4-Di-tert-butylphenol | -0.00 | 1.00 | 0.9564 | 0.9604 | 12.5 | 15.9 | ↓ |
| Glycine | -0.00 | 1.00 | 0.9604 | 0.9604 | 15.6 | 18.4 | ↓ |
