## Supplementary Table S2 for "β-Hydroxybutyrate elicits divergent metabolic responses between MCF-7 and T47D ER+ breast cancer cells under glucose restriction"

### Supplementary Table S2. TCGA Breast Invasive Carcinoma co-expression analysis (n=1,084 samples)

*Pairwise correlations between one-carbon/BCAA metabolism genes (SHMT1, SHMT2, BCAT1) and ketone body catabolic genes (OXCT1, ACAT1, BDH1). Source: cBioPortal, TCGA PanCancer Atlas, mRNA expression (RSEM, batch normalized, $\log_2$). Bonferroni correction for 9 simultaneous tests: α = 0.05/9 = 0.0056. The four pairs meeting both Bonferroni-corrected significance and a minimum correlation magnitude (|r| > 0.10) are highlighted.*

| Gene 1 | Gene 2 | Pearson r | Pearson p | Spearman ρ | Spearman p | Meets Bonferroni + \|r\| > 0.10 |
|---|---|---|---|---|---|---|
| SHMT1 | BDH1 | +0.18 | 2.88e-9 | +0.20 | 2.16e-11 | **Yes** |
| SHMT1 | OXCT1 | +0.03 | 0.4000 | −0.01 | 0.7000 | No |
| SHMT1 | ACAT1 | +0.05 | 0.1330 | +0.07 | 0.0312 | No |
| SHMT2 | BDH1 | +0.06 | 0.0606 | +0.07 | 0.0218 | No |
| SHMT2 | OXCT1 | +0.17 | 3.46e-8 | +0.13 | 1.17e-5 | **Yes** |
| SHMT2 | ACAT1 | −0.03 | 0.4000 | −0.03 | 0.3970 | No |
| BCAT1 | BDH1 | −0.36 | 3.69e-35 | −0.40 | 1.43e-43 | **Yes** |
| BCAT1 | OXCT1 | +0.16 | 1.10e-7 | +0.12 | 5.82e-5 | **Yes** |
| BCAT1 | ACAT1 | +0.03 | 0.2870 | +0.05 | 0.0712 | No |

*The strongest association is BCAT1–BDH1 (Spearman ρ = −0.40, $R^2$ = 0.13); the remaining three significant pairs are weak in magnitude ($R^2$ ≈ 0.03) and are interpreted as exploratory transcriptional associations rather than evidence of direct mechanistic coupling.*
