## Supplementary figures and images for "β-Hydroxybutyrate elicits divergent metabolic responses between MCF-7 and T47D ER+ breast cancer cells under glucose restriction"

### Supplementary Figure 1

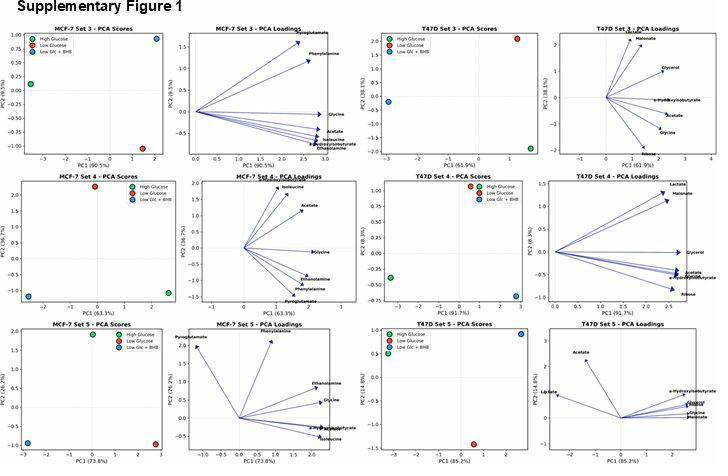
